# Tracking neurons across days with high-density probes

**DOI:** 10.1101/2023.10.12.562040

**Authors:** Enny H. van Beest, Célian Bimbard, Julie M. J. Fabre, Flóra Takács, Philip Coen, Anna Lebedeva, Kenneth Harris, Matteo Carandini

## Abstract

Neural activity spans multiple scales from milliseconds to months. Its evolution can be recorded with chronic electrodes, and especially with high-density arrays such as Neuropixels probes, which measure each spike at tens of sites and record hundreds of neurons. These arrays often record units with consistent spike waveforms over time, but produce vast amounts of data that require new approaches for tracking neurons across recordings. To meet this need, we developed UnitMatch, an open-access pipeline that operates after spike sorting, based only on each unit’s average spike waveform. We tested UnitMatch in Neuropixels recordings from the mouse brain, where it tracked neurons across weeks. In visual cortex, a neuron’s selectivity for visual stimuli and correlation with other neurons remained stable over days. In striatum, neuronal responses changed across days during learning of a task. UnitMatch is thus a promising tool to reveal invariance or plasticity in neural activity across days.

## Introduction

Neural activity spans a multitude of time scales, from the milliseconds that separate spikes to the days or months that characterize memory, learning, or aging. Changes at these longer timescales can be studied with two-photon imaging, where the same neurons can be visually tracked across days^1–5^. However, imaging methods miss the fast timescales and are hard to implement outside of cortex or in multiple areas at once. To cover all time scales across the brain, the ideal method is to use chronic electrodes.

Recordings with chronic electrodes reveal units (putative neurons) with consistent spike waveforms across days^6–20^. This constancy indicates that the units track the same neurons over time, particularly when the spikes are measured at multiple locations with stereotrodes^16^, tetrodes^14,15,19,21–26^, microwire bundles^27,28^, silicon probes^20,29^ and Neuropixels probes^30^.

In addition to spike waveform, the tracked units can maintain distinctive firing properties such as inter-spike interval distribution^10,11,13,20,27,28^, and functional properties such as sensory, cognitive, or motor correlates^11,14–16,23,27,28,30,31^. Consistent functional properties across days can provide a strong indication that a neuron has been tracked, but functional properties are not necessarily constant over time, and their variation is often the scientific question being investigated^6,7,20– 22,24,25,27,31–33.^

Tracking neurons across days has become more appealing with high-density arrays such as Neuropixels probes^30,34^. These probes are readily implanted chronically^30,35–38^, yielding hundreds of potentially matchable neurons across days. In addition, their geometry and density allow for reliable correction of electrode drift. All electrodes tend to drift relative to the brain, so that recorded neurons may appear and disappear, or change their apparent spike waveform. With rigid probes such as Neuropixels, however, most of the movement is likely to be parallel to the probe, causing the spikes to move from some recording sites to others. It is possible to compensate for this movement in software, so that neurons can be tracked robustly both within and across days^30,39^.

However, the current methods for matching neurons across days cannot process the vast amounts of data produced by multiple days of Neuropixels recordings. For example, one method relies on concatenating two recordings and spike sorting the resulting file^29,30^. This method is robust for pairs of recordings but becomes unwieldy for longer sequences: it does not scale to dozens of recordings obtained across weeks or months.

To solve this problem, we developed a pipeline called UnitMatch, which operates after spike sorting: each recording is spike sorted independently, without concatenation, with the user’s preferred algorithm. UnitMatch then deploys a fast and scalable naïve Bayes classifier on the units’ average spatiotemporal waveform in each recording, and tracks units across recordings assigning a probability to each match.

We tested UnitMatch in cortical and subcortical brain regions and found that it reliably tracked neurons across days and weeks. Its performance compares well to the concatenated method and to manual curation by human experts, while being both much faster and applicable to larger numbers of recordings.

Because UnitMatch does not rely on any properties of a unit beyond its spike waveform, it can be used to test whether these properties change over time. We used it to examine simple distinctive properties of neurons in mouse visual cortex, such as the correlation with other neurons or with visual stimuli. These properties remained remarkably stable, further confirming the ability of UnitMatch to track neurons across days. We also used UnitMatch to characterize the changes of neural representations in striatum neurons during learning, showing that UnitMatch can track neural dynamics in multiple brain regions on both fast and slow time scales.

## Methods and Results

UnitMatch tracks units across recordings that have been individually spike sorted (Figure 1). It takes as input the average spatiotemporal spike waveform of each unit in each half of each recording (Step 0). It then extracts key parameters from this waveform (Step 1) and uses them to compute similarity scores for each possible pair of units (Step 2). It performs within-day cross-validation to identify a similarity score threshold for putative matches (Step 3). It then corrects for drift (Step 4) across recordings and repeats Steps 1-3 to recompute the putative matches. Finally, it builds probability distributions for the similarity scores for putative matches and feeds them to a classifier to assign a probability to every possible match, suggesting unique identities to all units across recordings (Step 5).

**Figure 1.**
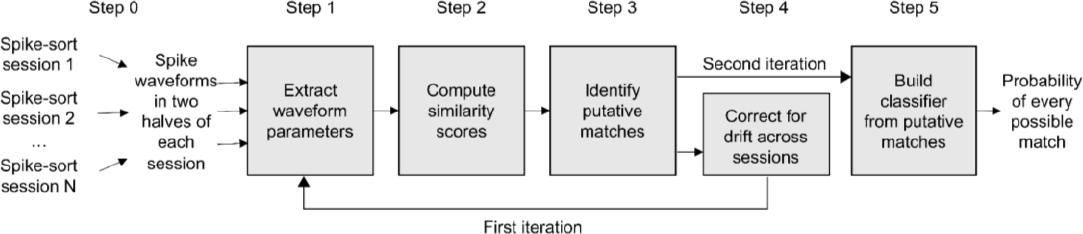
UnitMatch workflow. UnitMatch is an algorithm to find matches across multiple recording sessions. It operates after spike sorting (step 0), and it comprises of 5 steps done in two iterations.

Below, we describe these steps and illustrate them on a large body of data obtained in our laboratory. The description in the text is qualitative, and the equations are provided in Supplementary Information, section Mathematical definitions. We then assess the quality of matches by comparing the results with other approaches, and by measuring functional properties of the neurons that were remarkably consistent across days. Furthermore, we show that UnitMatch can track functional properties that change during learning.

### Step 0. Spike sorting

Before using UnitMatch, the users record neural activity in multiple sessions, and use their preferred software to spike sort each recording independently. For each recording, the results are put in a directory where each unit contributes a file with the average spatiotemporal spike waveform in the first and in the second half of the recording. These files have no information on individual spikes.

To develop, test, and illustrate UnitMatch, we used over 1,500 recording sessions performed over multiple days (up to 183) in mice implanted with chronic Neuropixels probes^30,34,38^ in multiple brain regions including cortex, hippocampus, striatum, and superior colliculus (Table S1). Each recording session was individually spike sorted with Kilosort^40^, which provides drift correction within each session^30^. After spike sorting, we used a set of quality measures^41^ to select 21.9 ± 11.5% (mean ± s.d. n = 1,514 recording sessions across 17 mice) units that were well isolated and distinct from noise (Figure S1).

### Step 1: Waveform parameters

High-density recording arrays such as Neuropixels probes (Figure 2A) sample the spikes of an individual unit at many recording sites, revealing the unit’s characteristic spatiotemporal waveform (Figure 2B). The amplitude of the waveform peaks at a maximum site and decays gradually with distance from that site (Figure 2B,C). UnitMatch fits this decay with an exponential function and obtains the distance *d*_10_at which the amplitude reaches 10% of the maximum (Figure 2C). In the example recordings, this value ranged between 30 and 114 μm (95% confidence interval, Figure 2D). For each unit, UnitMatch considers the recording sites closer than *d*_10_(but at most 150 μm away) from the maximum site. In our data, this typically resulted in 6-24 sites arranged in two columns (e.g., Figure 2A-B).

**Figure 2.**
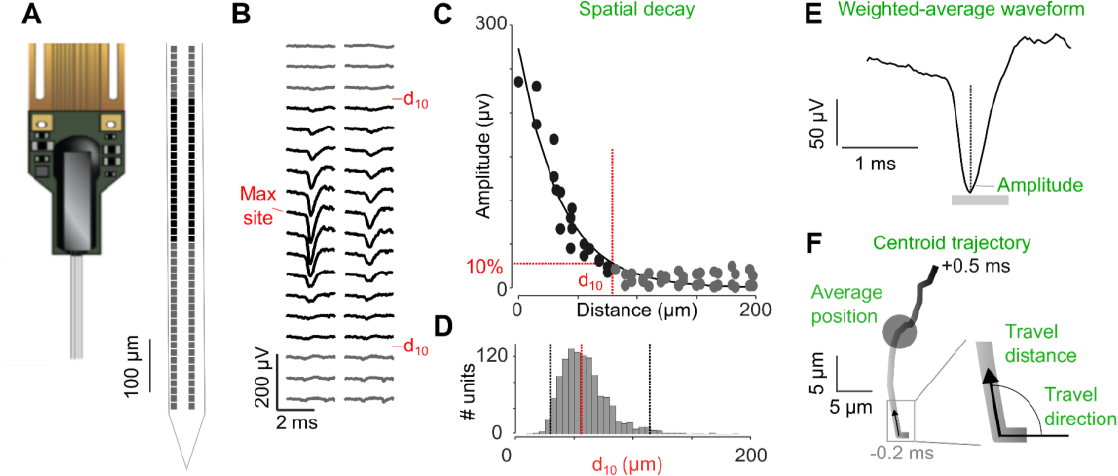
Extracting spike waveform parameters. (A)A 4-shank Neuropixels 2.0 probe (*left*), showing the bottom of one shank (*right*) and its recording sites (*squares*). (B)Average spike waveform for an example unit, in the 24 recording sites marked in *black* in A and in 12 adjacent sites (*gray*). (C)Amplitude of the waveform as a function of distance to ‘max site’ for the example unit. Using an exponential decay fit (*curve*), we defined the distance d_10_ at which the amplitude drops to 10% of the maximum. <u>Spatial decay</u> is computed from the slope of the amplitude decrease over distance. (D)Distribution of d_10_ for all units in two example recordings, showing the median (*dashed red line*) and the 95% confidence interval (*dashed black lines*). (E)<u>Weighted-average waveform</u> for the example unit in B, C, computed by giving larger weight to sites near the maximum site. The unit’s amplitude is taken from this waveform. (F)<u>Centroid trajectory</u> of the example waveform from 0.2 ms before the peak (*bottom*) to 0.5 ms after the peak (*top*), showing the <u>average centroid</u> (*circle*). <u>Travel direction and distance</u> are calculated at each time point.

For each unit, UnitMatch uses the spatiotemporal spike waveform measured at the selected recording sites to extract seven attributes:

- The spatial **decay**, i.e., the rate at which amplitude decreases with distance from the maximum site (Figure 2C, Equation 8).
- The weighted-average **waveform** (Figure 2E; Equation 9) obtained by averaging
- waveforms across sites, weighted by the maximum amplitude of each site.
- The **amplitude** of that weighted-average waveform (Figure 2E, Equation 10).
- The average **centroid** (Figure 2F, Equation 6), defined as the average position weighted by the maximum amplitude on each recording site.
- The **trajectory** of the spatial centroid from 0.2 ms before the peak to 0.5 ms after the peak (Figure 2F, Equation 4).
- The **distance** travelled at each time point (Figure 2F).
- The travel **direction** of the spatial centroid at each time point (Figure 2F, Equation 5).

### Step 2: Similarity scores

After extracting the spatiotemporal waveform parameters, UnitMatch compares these parameters for every pair of waveforms within and across days, computing six similarity scores using the extracted parameters:

- **Decay** similarity (*D*; Equation 14); spatial *decay*.
- **Waveform** similarity (*W*, Equation 18); *waveform* correlation and normalized difference averaged.
- **Amplitude** similarity (*A*; Equation 13)
- **Centroid** similarity (*C*, Equation 20)
- **Volatility** similarity (*V*, Equation 23); captures the stability of the difference between *centroids* over time.
- **Route** similarity (*R*; Equation 24): captures the overall similarity of the *trajectory*: *direction* and *distance* travelled.

Each similarity score is scaled between zero and one, with one indicating the highest similarity. Finally, in addition to the six individual scores, we also average the scores to compute a **total** similarity score *T*.

To gain an intuition for these scores, consider their values for two example pairs of units. The first example involves two neighboring units recorded on the same day (Figure 3A,B). Because they are near each other, they have high centroid similarity *C*. However, their spike waveforms are not similar (low value of *W*), and so are their spatial decays (low value of *D*) and routes (low value of *R*). As a result, the total similarity score *T* is well below 1 (Figure 3C). Conversely, in the case of a single unit recorded in two different days we observed similar waveforms and trajectories (Figure 3D,E), with high values of most similarity scores and consequently a total similarity score *T* near the maximal value of 1 (Figure 3F).

**Figure 3.**
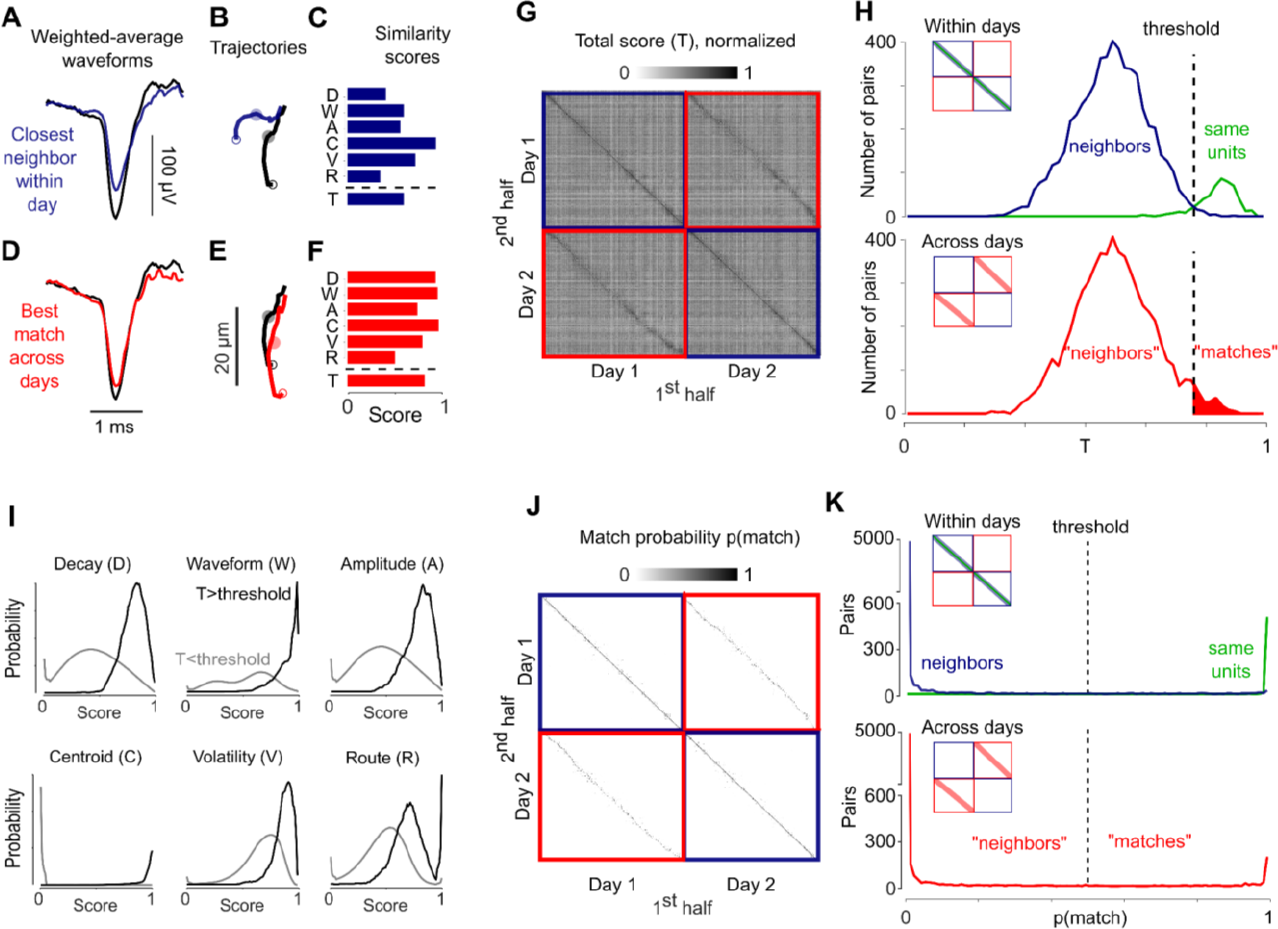
Computing similarity scores and setting up the classifier. (A) Average weighted waveform for the example unit in Figure 2 (*black*) and for a different unit (the nearest physical neighbor) from the same recording (*blue*). (B) Centroid trajectories for the two units. (C) The six similarity scores between the two units and their average, the total similarity score *T*. (D-F) Same as (A-C), comparing the example unit (*black*) with the most similar unit across days (*red*), which was very likely the same neuron. (G) Total similarity score for all pairs of units within days (*blue squares*) and across days (*red squares*), for an example pair of recordings, showing the 1^st^ half of each recording (*columns*) vs. the second half (*rows*). (H) *Top*: Distribution of total similarity scores in the two halves of a recording day, showing for the same units the two halves (*green*) and for other neighboring units (*blue*; centroid <50 μm away). *Bottom*: same, for units measured across days (*red*) after drift correction. The threshold for putative matching (*dashed line*) depends on the number of units and of recordings. (I)Probability densities of each similarity score, for putative matches (*black*) and for putative non-matches (*gray*). (J) Match probability computed by the naïve Bayes classifier trained with the probability distributions in (I). Format as in G. (K) *Top*: Distribution of match probabilities within two halves of the same day for same units (*green*) and neighbors (*blue*). Format as in H. *Bottom* same, for units recorded across days (*red*). If probability is >0.5, UnitMatch defines a pair as a match.

### Step 3: Putative matches

We can now measure the total similarity score *T* for each pair of units within and across days. *T* tended to be high when applied to the same unit recorded across the two halves of a recording (main diagonal in Figure 3G).

To identify putative matches, we defined a threshold as the value of *T* above which there were more pairs of waveforms from the same unit (green curve in Figure 3H, *top*) than pairs of waveforms from neighboring units (average centroid distance <50 μm, blue curve in Figure 3H, *top*) (Equation 28).

The distribution of *T* across days resembled the neighbor distribution measured within days (red curve in Figure 3H, *bottom*), with larger tail above the data-driven threshold. We thus select pairs with values of *T* beyond this threshold as putative matches across days.

### Step 4: Drift correction

First, we equalized the means of the distributions within and across days, to account for overall lower scores across days prior to drift correction. Then, we computed the median centroid displacement of the population of the putative matches across days. After applying this rigid transformation to all parameters affected by position, we then recalculated all the values computed in the previous three steps, thus finding a more robust set of putative matches. Results shown in Figure 3 are after this drift correction. Note that within-recording drift correction is taken care of by the spike sorting algorithm we used.

### Step 5: Match probabilities

Having used the total similarity score *T* to identify putative matches (Figure 3H), UnitMatch then goes back to the individual similarity scores and uses their distributions to train a classifier. The total similarity score identifies putative matches (*T*>threshold) and putative nonmatches (*T*<threshold, Figure 3H). Plotting the distributions of the six similarity scores for these pairs reveals major differences (Figure 3I). Based on these distributions, we defined a naïve Bayes classifier (Equation 29), which takes as input the values of the six similarity scores for two spike waveforms and outputs the ‘match probability’: the posterior probability of the two waveforms coming from the same unit.

This classifier correctly identified matches within a day and indicated that some pairs of units had a high probability of match across days. Within a day, matching probabilities were close to 1 for the same units measured in the first and second half of the recording (main diagonal in Figure 3J, *green curve* in Figure 3K, *top*), and were close to zero for neighboring units (*blue* in Figure 3K, *top*). Across days, most pairs of unit waveforms are also expected to come from different neurons, which is reflected in a large fraction of matching probabilities close to 0 (Figure 3K, *bottom*). However, a small proportion of unit pairs across days had a matching probability close to 1. These matches reflect units tracked across days.

### Performance metrics

We first evaluated UnitMatch’s performance using the cross-validation of the units’ waveforms within days, and found low percentages of false positives and false negatives (Supplementary analyses, Figure S3A). From the maximum possible number of units recorded across two consecutive days, UnitMatch found 31.3 ± 11.2% (median ± m.a.d., n = 446 pairs of days) to be a match. Reassuringly, when we applied UnitMatch to acute recordings, where the probe was reinserted daily and had negligible chance of finding the same unit, finding a match was rare (4.0 ± 2.8%, n = 18 pairs of consecutive days; Wilcoxon rank sum comparing chronic and acute: p < 10^-10^; Figure S3B). Note that this was when UnitMatch was pushed to find matches by pretending the probe was in the same location between the recording sessions.

Next, we compared the performance of UnitMatch to expert manual curation and to spike sorting performed on stitched recordings^29,30^. We found that UnitMatch performed more similarly to manual curation than to the sorting on stitched recordings (Supplementary analysis, Figure S3CD). Sorting the stitched recordings with Kilosort tended to overestimate the number of matches across recordings, specifically for noisier datasets. The higher similarity between UnitMatch and manual curation is reassuring because the latter is generally regarded more highly than automated spike sorting. However, neither can be considered ground truth.

### Validation with stable functional properties

A more reliable estimate can be found by assessing the stability of the neurons’ functional activity. If this activity is found to be distinctive across neurons and stable across days, one can conclude that the tracking algorithm performed well. However, if the activity is found to change over days, then one cannot make such an inference. In our case, we found functional activity to be remarkably stable, yet distinctive across neurons, and thus could use this stability to validate UnitMatch performance.

We thus looked for functional ‘fingerprints’, i.e., patterns of activity that are known to be distinctive and are potentially stable across days. We considered three such fingerprints: a unit’s autocorrelogram (the correlation of the unit’s firing rate with its own firing rate at nearby times^8,10,11,13,20,23,27,28^), a unit’s response to a large set of visual stimuli (when the unit was in visual cortex^27,28,30,31^), and a unit’s population coupling (the correlation of its firing rate with the other units recorded at the same time^11,23,42,43^).

The autocorrelogram (ACG) of tracked units remained highly consistent across days. It has been used as a feature to track units across days^10,11,13,20,27,28^ or as a diagnostic of this tracking^8,23^. Accordingly, the ACGs were typically different for neighboring units but similar for units matched across days (Figure 4A), which are two properties that are necessary for a good fingerprint. In an example mouse, we observed that the distribution of correlation for a pair of ACGs coming from matches across days was high (Figure 4B), and to the same extent as cross-validated within-day comparisons. On the contrary, different units had much lower ACG correlations.

**Figure 4.**
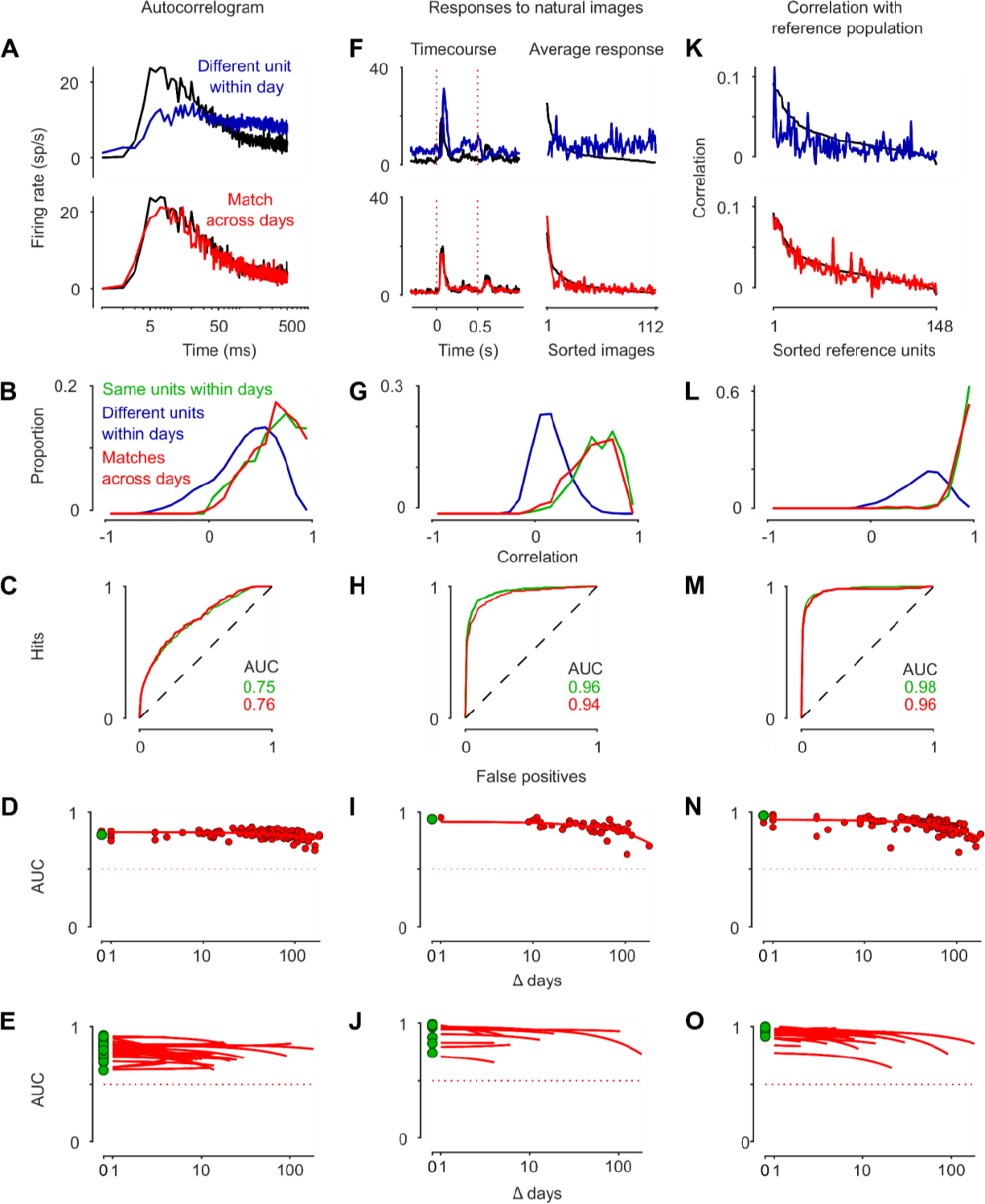
Validation with stable functional properties. (A)Autocorrelogram (ACG) of an example a unit and one of its neighbors (*top*) or its match on the next day (*bottom*). (B)Distribution of the ACGs correlation for pairs of waveforms coming from the same unit (*green*) or different units (*blue*) within days and matches across days (*red*). Data from two consecutive days in an example mouse. (C)Receiver operating characteristic (ROC) curves when classifying the ACGs correlations of the same vs. different units within days (*green*) or matching versus non-matching units across days (*red*). Data from two consecutive days in an example mouse. (D)Area under the curve (AUC) for many pairs of days, spaced by different amounts (*red dots*). The AUC of the same vs. different units within recordings is also shown (*green*). Stability was estimated with a linear fit (*curve*). The discriminability remained highly stable, even across months. Data from an example mouse. (E)AUC for many pairs of days, across many mice and recording locations. (F)Comparison of the responses to natural images for a unit and one of its neighbors (*top*) or its match on the next day (*bottom*). The responses to the images were summarized by averaging over all images to obtain the time course (*left*) or averaging over time to obtain the responses to individual images (*right*). These two profiles were then concatenated to form a single fingerprint, which was compared across units. Images were ordered by decreasing response of the black unit. (G-J) Same as (B-E) but using the responses to natural images as a fingerprint. (K) Correlation of the firing rate of a unit with other units forming a reference population that was tracked across days using UnitMatch. The correlation of a unit (*black*), one of its neighbors (*top, blue*) or its match on the next day (*bottom, red*) is shown. The neurons in the reference population were ordered by decreasing correlation with the unit from day 1 (*black*). (L-O) Same as (B-E) but using the correlation with a reference population as a fingerprint.

We quantified the separation of the distributions of the correlations of matched and non-matched pairs by computing the receiver operating characteristic (ROC) curve (Figure 4C) and quantified the discriminability index as the area under the ROC curve (AUC). The AUC was high for this example pair of days (0.76), suggesting that the value of the ACGs correlation for a pair was predictive of whether it was a match or not. This held true even when increasing the number of days between recordings (Figure 4D), and across all mice (Figure 4E). On average, the AUC was 0.78 ± 0.01 across days (0.80 ± 0.01 within days, mean ± s.e., n = 17 mice) and decayed slowly with each additional day between recordings (-0.003 ± 0.049, median ± m.a.d., n = 17 mice). For the example mouse, the AUC remained at 0.78 after 183 days.

Units in visual cortex that were tracked across two days also typically showed consistent responses to visual stimuli. Neurons in mouse visual cortex give distinctive responses to natural images, and these responses are reproduced across days^27,28,30,31^. Consistent with this, a typical “match” returned by UnitMatch across days gave similar responses to natural images on each day (Figure 4F). This set of responses provides a signature that was consistent both within and across days. In the example mouse, it yielded AUCs of 0.94 across days vs. 0.96 within days (Figure 4G-H). Similar results were seen across mice, with AUCs of 0.86 ± 0.02 vs. 0.90 ± 0.02 (mean ± s.e., n = 10 mice). This held true even with long intervals between recordings (slope of -0.002 ± 0.009, median ± m.a.d., n = 10 mice, Figure 4I-J). For the example mouse, the AUC was still 0.70 after 183 days.

Finally, the population coupling of tracked units was also highly consistent across days. The population coupling is the instantaneous correlation of a unit’s activity with the activity of other neurons in the population^11,23,42,43^. The cross-correlation of spikes of a unit to a population of reference units (tracked across the two days) was highly consistent across days (Figure 4K). This cross-correlation with a reference population provided a distinctive ‘fingerprint’ that was highly correlated both within and across days (Figure 4L), and not for neighboring units. The discriminability of this measure was particularly high, with AUC indices close to 1 (0.96 across days vs. 0.98 within days), indicating that the pairs found by UnitMatch were indeed highly likely to be the same across days (Figure 4M). Again, this held true across mice (0.92 ± 0.01 vs. 0.96 ± 0.01 across mice, mean ± s.e., n = 17 mice) and even across weeks and months (-0.006 ± 0.030, median ± m.a.d., n = 17 mice), suggesting that the correlation patterns of the population of neurons were highly stable over time (Figure 4N,O). For the example mouse, the AUC was still 0.80 after 183 days. This fingerprint, along with the ACG, can be used in any region of the brain since it does not depend on the area’s responses to external stimuli.

Using the stability of functional properties, we once more compared the performance of UnitMatch to running our spike sorting algorithm on stitched recordings. This time we ran both algorithms on four concatenated recording sessions and evaluated their accuracy using functional measures. In line with the earlier curation results, there was a larger overlap between functional response stability and UnitMatch than with spike sorting the concatenated data. This was especially the case with recording sessions further apart (Figure S3E-F).

### Tracking units across learning

We have shown that UnitMatch can be used to track units across days, and this can be validated by stable functional scores. An important application of this algorithm is to track units as their functional responses evolve over time, particularly as a result of learning. To illustrate its potential, we applied UnitMatch to an exploratory dataset captured during a learning process.

We trained a mouse (Table S1) in a visuomotor operant task and recorded neural activity in the dorsomedial striatum using a chronically implanted Neuropixels probe. After implantation, the mouse was water-restricted and head-fixed in front of three screens with its forelimbs resting on a steering wheel. When the stimulus appeared on the left screen, moving the wheel clockwise moved the stimulus to the center, resulting in a sucrose water reward (Figure 5A). After each training day, we recorded passive responses to the presentation of the center stimulus. The mouse learned to correctly move the wheel over a two-day training period (Figure 5B).

**Figure 5.**
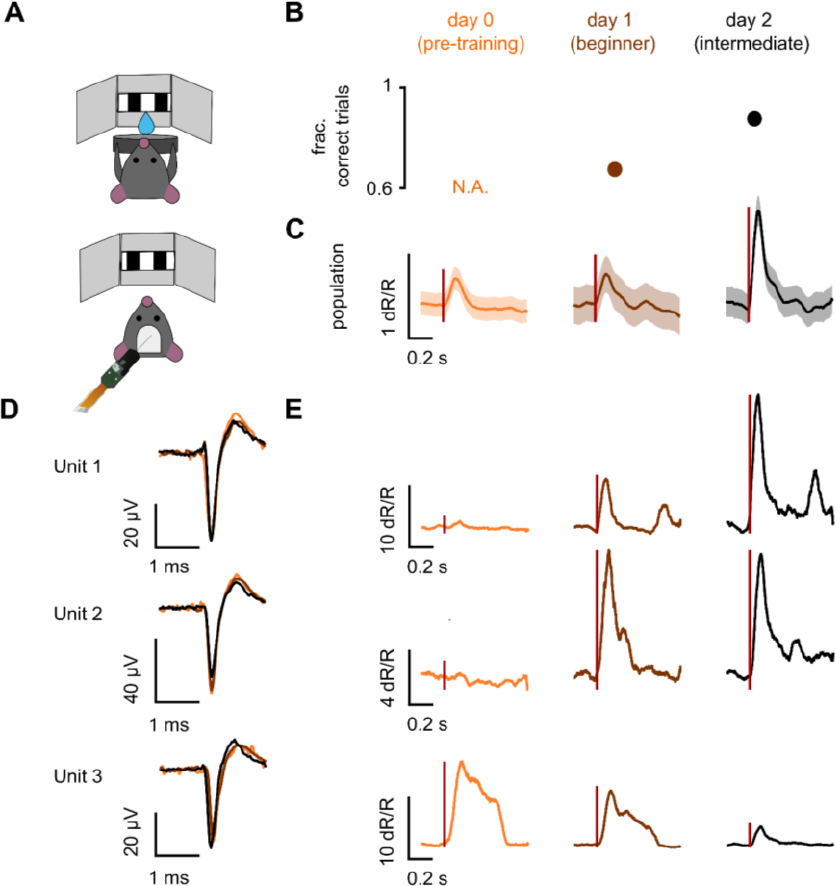
Tracking units across learning. (A)Task trial structure. (B)Fraction of correct trials as a function of training day (n=1 mouse). (C)Average baseline (-0.2-0 s before stimulus onset) normalized firing rate (dR/R) aligned to stimulus onset during passive viewing, sorted by training day (left: day 0, middle: day 1, right: day 2) for average population. Red line indicates stimulus onset. (D)Mean raw waveform for each training day for 3 example units (top: unit 1, middle: unit 2, and bottom: unit 3). (E)Same as C) but for three example units tracked across days by UnitMatch.

We analyzed data recorded during passive viewing of the same set of stimuli on day 0 (pre-training), day 1 (after the first training session), and day 2 (after the second training session). Without tracking neurons, it is possible to demonstrate that responses to the target stimulus increase over learning (Figure 5C). However, using the UnitMatch pipeline as described here, we were able to track individual units over multiple recording days (example units in Figure 5D). We found that responses of individual units to the target stimulus were highly variable. For example, some units (e.g., units 1 and 2) would not respond to the target stimulus prior to training, but developed a visual response with learning, either gradually (unit 1), or abruptly (unit 2). Conversely, other units (e.g., unit 3) initially had a strong visual response to the target, which decreased with learning.

This proof of concept shows that UnitMatch is a promising tool to reveal invariance or plasticity in neural activity across days.

## Discussion

We have shown that UnitMatch can track populations of neurons across days in data obtained with high-density arrays such as Neuropixels probes. UnitMatch is highly scalable because it operates after spike sorting, and because it makes few assumptions: it uses only the average spike waveform of each unit, without knowledge of its spike train. Additionally, rather than giving a binary output, UnitMatch provides a matching probability. Our tests with Neuropixels recordings from various mouse brain regions indicate that it successfully tracks neurons across weeks, and that it compares favorably to manual curation and to spike sorting on concatenated sessions.

UnitMatch revealed that distinctive functional properties of each neuron – autocorrelation, responses to visual stimuli, and population coupling – remain remarkably stable over days. Because these properties are highly distinctive across neurons, their stability is not only of interest by itself, but also indicates that UnitMatch indeed finds the same neurons across days. The population coupling fingerprint, especially, was highly accurate and could be used across various brain areas, without the need of any external stimuli. Using these functional metrics, we showed that UnitMatch performed better than spike sorting on concatenated recordings on long time scales.

Since functional properties of neurons are so stable over time, it is tempting to use functional properties themselves to track neurons. However, this would prevent any evaluation of the variation in functional properties across time, and such a variation has been documented^4,5,20^. For example, the slow decrease in AUC values that we observed across days could be explained either by a decrease in the quality of matching, or changes in functional properties of the units. Therefore, unless there is reason to believe that the functional properties are constant^9,17,31^, it is prudent to exclude these properties from the criteria that determine the tracking of units, and only consider them as a possible validation^11,15,16,19,30^ or as a separate question^6,7,21,22,25,31,32^. UnitMatch operates exclusively on the units’ waveforms and thus avoids circularity when addressing such questions. To illustrate this, we also tracked units that changed their functional responses over learning.

Though UnitMatch could track the same units over months, the number of units that were tracked declined with time. This decline could derive from numerous sources independent of the algorithm, such as a decline in recording quality, accumulation of uncorrected drift across recordings, neurons becoming silent or dying, or changes in waveform properties. Indeed, tracking of units was more stable when recordings were more stable, suggesting recording quality and drift are partially responsible (Figure S5). Ideally, further work will reveal the contribution of each of these factors to the quality of the tracking.

Taken together, these findings show that chronic high-density recordings and UnitMatch combined are a powerful tool to characterize neural activity spanning a multitude of brain regions and time scales, such as memory, learning, and aging.

## Acknowledgments

We thank Magdalena Robacha for help with experiments and histology, Bex Terry for animal husbandry, and Charu Bai Reddy for lab management. This work was funded by Wellcome Trust (Investigator Award 223144 to MC and KDH), Biotechnology and Biological Sciences Research Council (Grant BB/T016639/1 to MC and PC), European Council (Marie Skłodowska fellowship 101022757 to EvB), and European Molecular Biology Organization (ALTF 740-2019 to CB). MC holds the GlaxoSmithKline / Fight for Sight Chair in Visual Neuroscience.

## Code availability

UnitMatch software is published via Zenodo (DOI: 10.5281/zenodo.8435312) and available via GitHub (https://github.com/EnnyvanBeest/UnitMatch/tree/v1.0.0_UnitMatch).

## CRediT contributions

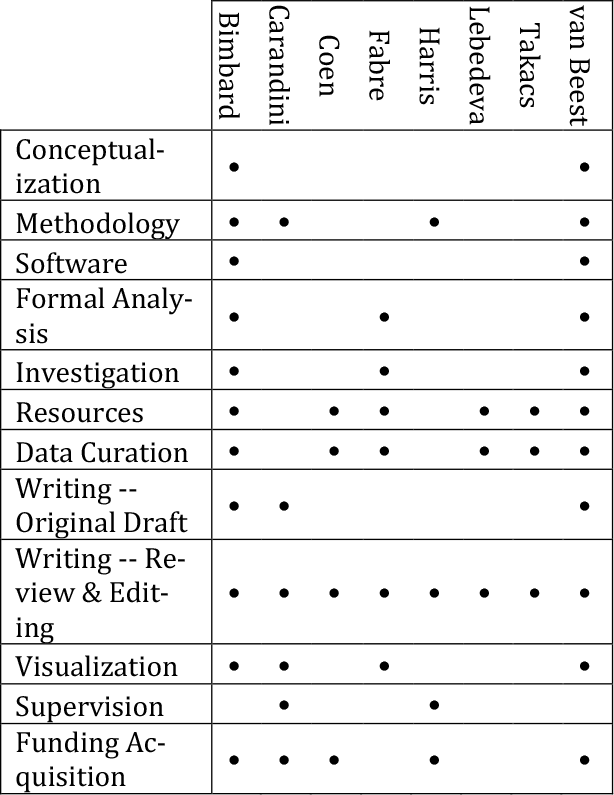

## Experimental methods

Experimental procedures were conducted at UCL according to the UK Animals Scientific Procedures Act (1986) and under personal and project licenses released by the Home Office following appropriate ethics review.

### Mice

We analyzed the data from 18 chronically implanted mice. During the experiments, mice were typically head fixed and exposed to sensory stimuli, engaged in a task, or resting. The mice were recorded from different experimental rigs, implanted, and recorded by different experimenters using different devices (Table S1).

### Surgeries and implants

#### Headplate surgery

A brief (around 1 h) initial surgery was performed under isoflurane (1–3% in O_2_) anesthesia to implant either a titanium headplate (∼ 25 × 3 × 0.5 mm, 0.2 g in the case of the Apollo implant) or a steel headplate (∼ 25 × 5 × 1 mm, xx g in the case of the Ultralight and Cemented implants). In brief, the dorsal surface of the skull was cleared of skin and periosteum. A thin layer of cyanoacrylate was applied to the skull and allowed to dry. Thin layers of UV-curing optical glue (Norland Optical Adhesives #81, Norland Products) were applied and cured until the exposed skull was covered. The head plate was attached to the skull over the interparietal bone with Super-Bond polymer. In one mouse (ID 2/19), a silver wire was implanted in the mouse’s skull in order to ground the mouse during recordings. After recovery, mice were treated with carprofen or meloxicam for three days, then acclimated to handling and head-fixation. Mice were then implanted with either a modular recoverable^38^, Ultralight or cemented implant (see below). Briefly, craniotomies were performed on the day of the implantation, under isoflurane (1–3% in O2) anesthesia, and after injection of Colvasone and Rimadyl. The UV glue was removed, and the skull cleaned and scarred for best adhesion of the cement. The skull was levelled, before opening the craniotomies using a drill or a biopsy punch. Once exposed, the brain was then covered with Dura-Gel (Cambridge Neurotech).

#### Cemented implant

The data from four animals was already published^30^. In short, these four animals were implanted by holding and inserting the probes using a cemented dovetail and applying dental cement to encase the probe PCB and reliably attach it to the skull. The recordings were made in external reference mode, using the Ag wire or the headplate as the reference signal.

#### Recoverable modular implants

Three different recoverable, modular implants were used. The methods for the Apollo implant^38^ and the ‘Haesler’ implant^30,37^ have been described in their respective papers. The third implant (‘Isogai’ implant) is conceptually similar. In short, the implant was held using the 3D-printed payload holder and positioned using a micromanipulator (Sensapex). After carefully positioning of the shanks at the surface of the brain, avoiding blood vessels, probes were inserted at slow speed (3-5 μm/s). Prior to surgery, the probes were coated with DiI by either manually brushing each probe with a droplet of DiI or dipping them in directly in DiI, for histological reconstruction. Once the desired depth was reached (optimally just before the docking module touched the skull), the implant was sealed using UV glue, then covered with Super-Bond polymer, ensuring that only the docking module was cemented. After finishing all recording sessions, the probes were explanted and cleaned before reusing. The recordings were made in external or internal reference mode, using the headplate as the reference signal.

#### Ultralight implant

To record chronically while a mouse (ID 19) learned a task and up to 3 probes simultaneously, we developed an ultralight implant (https://github.com/Julie-Fabre/ultralight_implant). Briefly, an implant was constructed using between 1 to 3 Neuropixels probes encased in rigid-resin K custom-made 3D printed parts. A thin square of sorbuthane sheet was added to the front of the implant. Special care was taken to ensure all shanks were parallel to each other and to the implant. This implant was then slowly lowered into the brain. At the target depth, the implant base was covered in vaseline to protect the shanks from subsequent cement applications. We then applied cement to the implant and mouse skull. To explant, we carefully drilled the implant out in the areas where Vaseline had been applied.

#### Data processing

Electrophysiology data were acquired using SpikeGLX (**https://billkarsh.github.io/SpikeGLX/**) and each session was spike sorted with Kilosort^40^ (Table S1). Data were preprocessed using ‘*ExtractKilosortData*.*m’*, meaning that all relevant information was extracted (e.g., positions of recording sites, information on extracted clusters and their spike times) and common noise was removed. Well-isolated units were selected using Bombcell (**https://github.com/Julie-Fabre/bombcell**, using parameters defined in bc_qualityParamValuesForUnitMatch.m). For each session the average waveform on every recording site for each unit was extracted, either through Bombcell or through Unitmatch’s *‘ExtractAndSaveAverageWaveforms*.*m’*.

Input to the core of UnitMatch, which is matching units purely based on waveforms, was information on the clusters, at least 1) cluster identity, 2) a Boolean on which clusters to include – typically well isolated units, 3) which recording session it was recorded in, and 4) on which probe it was recorded. Additionally, it requires parameters (we used default parameters available using *‘DefaultParametersUnitMatch*.*m’*), containing information on where to find the raw waveforms.

Example analysis pipelines from raw electrophysiology recorded using SpikeGLX all the way to using and validating UnitMatch are provided in the UnitMatch repository. A minimal use case scenario is also provided in *‘DEMO_UNITMATCH*.*m’*, which is also useful for electrophysiological data recorded and preprocessed using other probes and software.

## Supplementary Information

### Mathematical definitions

We consider recordings made in a probe with *N* sites, and we denote with *p*_*s*_ the position of site *s* (a vector with the x,y coordinates). For every neuron *i* we denote the spike waveform at site *s* and at time *t* as *w*_*s,t,i*_ (averaged across *n* spikes of that neuron).

### Step 1: Waveform parameters

Some useful summaries of the spike waveform include the spatial footprint:

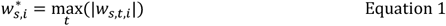

and the maximum site 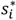 where the voltage has maximum amplitude:

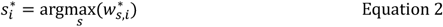

Most analyses are performed in a time window of size *T* samples starting 0.23 ms before the waveform reaches its peak and ending 0.50 ms after the peak. To establish a baseline noise level, we used a window of same duration starting 1.33 ms before waveform onset.

The spatial decay of the waveform is the degree to which the waveform’s maximum amplitude at site *s* decreases as a function of distance from the peak site, 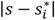. To describe it, we fit an exponential decay function (Figure 1C) with scale λ_*i*_ such that:

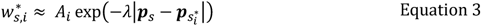

and we use this fit to obtain the distance at which the amplitude drops to 10% of maximal value, *d*_10_= log(10)/λ. For further analysis we only take recording sites into account with distance to 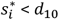.

The centroid trajectory of neuron *i* is (Figure 1F):

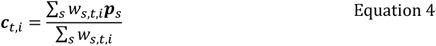

and its travel direction at each time *t* is:

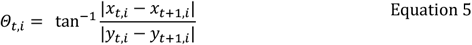

*x*_*t,i*_ and *y*_*t,i*_ being the components of ***c***_*t,i*_.

The neuron’s average centroid (Figure 1F) is:

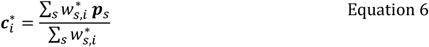

To calculate a neuron’s average waveform, we start by computing the proximity *f*_*s,i*_ of each site *s* to the centroid of the neuron 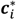

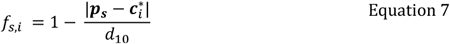

where *d*_10_is the distance where amplitude drops to 10% (or 150 μm if that distance is larger). At sites that are further away (were *f*_*s,i*_ would be negative) we set *f*_*s,i*_ =0.

We then calculate the unit’s spatial decay as the average decrease in amplitude divided by increase in distance for all sites closer than *d*_10_(Figure 2)

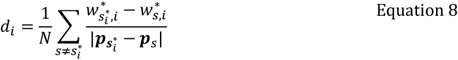

We then compute the neuron’s weighted-average waveform 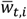 (Figure 2E) as

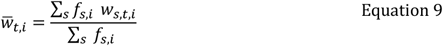

We use this waveform to compute the weighted amplitude of the neuron’s spike:

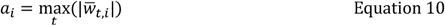

When comparing waveforms between units we normalize 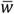 to obtain

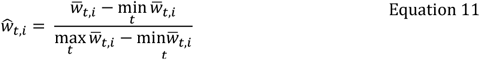

### Step 2: Similarity scores

Based on these parameters, we next compute similarity scores for each pair of units *i* and *j*. These scores are scaled between 0 and 1, with 1 being the most similar. For most similarity scores, we do “0-99 scaling”: we rescale the similarity scores so that the minimum is 0 and the 99^th^ percentile is 1. If *X*_*i,j*_ is the similarity score between units *i* and *j*, its 0-99 scaling is:

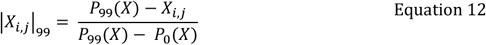

where *P*_*K*_(*X*) is the K-th percentile of X. For similarity scores above the 99^th^ percentile, we clip the score to 1.

We used two types of similarity scores: those based waveform timecourses and those based on waveform trajectories.

#### Amplitude similarity

We compute the difference in maximum amplitude between each unit *i* and *j*, and we apply 0-99 scaling to its square root:

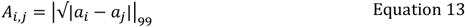

#### Decay similarity

We compute the difference in spatial decay and we apply 0-99 scaling to it:

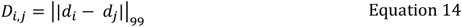

#### Waveform similarity

We compute the Euclidean distance between the waveforms, and we apply 0-99 scaling to it:

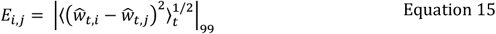

We also compute the correlation between the waveforms and apply Fisher’s z-transformation and 0-99 scaling to it:

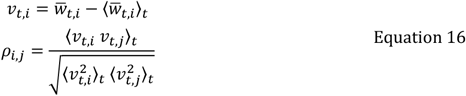

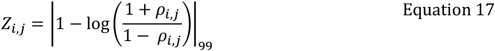

Empirically, we found both measures (distance and correlation) to be informative. Of course, they are also highly correlated with each other (Figure S2B). This correlation poses problems for a naïve Bayes decoder. To take them both into consideration, we defined ‘waveform similarity’ as their average.

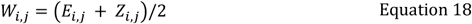

#### Centroid similarity

We compute the mean absolute distance between two centroids and then we rescale it to obtain a measure of proximity that is 1 if centroids are identical and 0 if they are further than *d*_*max*_= 100 μm:

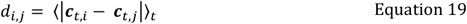

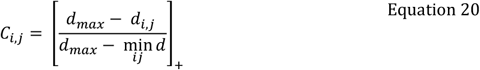

Units that are further apart than *cd*_*max*_are unlikely to be a match, even when considering drift between recordings.

#### Volatility similarity

If some of the drift remains uncorrected, a unit that appears in two recordings may have centroid trajectories that are identical but displaced by a constant shift. To correct for this, we subtracted the average centroid (Equation 6) from the centroid trajectory (Equation 4) for each unit and computed their similarity *F*_*i,j*_ across units as in Equation 19 and Equation 20.

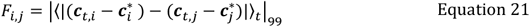

We also compute the standard deviation in Euclidean distance between centroids, and apply 0-99 scaling to it:

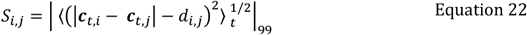

Since *F*_*i,j*_ and *S*_*i,j*_ are highly correlated (Figure S2B), we averaged these two scorers to centroid ‘volatility’ similarity:

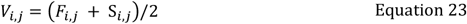

#### Route similarity

We compute the summed difference in direction (angle) of the centroid trajectory, and apply 0-99 scaling to it:

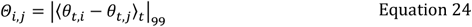

Additionally, we compute the distance travelled by the centroid between each time point of the trajectory and compare the differences between each pair of units *i* and *j*, and apply 0-99 scaling.

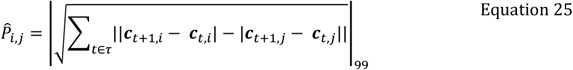

The final route similarity

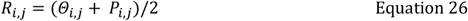

#### Default similarity scores

Before settling on this set of default similarity scores, we evaluated the performance of other scores (Figure S2). For each set of scores, we computed the AUC value in classifying whether two waveforms came from the same unit or not (Figure S2A). This process led us to consolidate similarity scores that were highly correlated with each other (Figure S2B). Note that, based on within day cross-validated performance, a user of UnitMatch will be advised what similarity scores to use for every individual dataset. In this paper, we only used default parameters and scores.

### Step 3 and 5: Identifying matches

Having defined these six similarity scores for each pair of units *i* and *j*, we averaged them to obtain a total score:

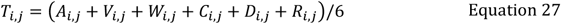

We define preliminary class (*M*) between two units as:

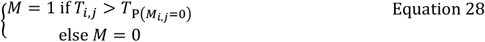

In which *T*_p(*Mi,j*=0)_ is defined by finding where the probability distributions of *T*_*i,i*_ and *T*_*i≠j*_ (neighbors within days) cross. In the case of overall lower scores across days (e.g. due to uncorrected drift) we lowered the threshold by the difference in means (by fitting a normal distribution) for the within day distribution (blue and green curves combined in Figure 3H, *top*) and the across days distribution (red curve in Figure 3H, *bottom*).

We use the preliminary class labels to build the probability distributions for the similarity scores as defined above, and use these to compute the probability of a match between units *i* and *j* as

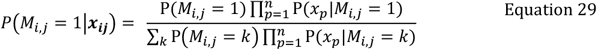

### Functional scores

To evaluate UnitMatch performance, we determined various functional scores of neuronal activity.

### Autocorrelogram fingerprint

For each neuron *i* with spike train *T*_*i,t*_ we compute the autocorrelogram *A*_*i*_ of elements *a*_*i,s*_ as the average number of spikes/seconds around a spike at defined time intervals *τ* (1ms bins, 1 second duration centered on the spike).

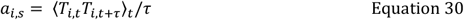

The autocorrelogram *A*_*i*_ was then use as the first functional fingerprint.

### Natural image responses fingerprint

To characterize the functional responses of the neurons in visual cortex, we showed 112 natural images, each presented 5 times in a random order, to the head-fixed mice ^30^. Two versions of the protocol were used, one long (1s-stimulus, 2s-intertrial interval), and one short (0.5s, 0.8s), without affecting the overall reliability of the fingerprints. To define the fingerprint, we computed the responses as the peristimulus histograms locked on the image onset (0.3s before and 0.5s after) and the image offset (from 0 to 0.5s after), using 5ms bins. The response *R*_*i,t,s*_ for each unit *i* and stimulus *s* were then defined as the concatenation of the onset and offset matrices along their temporal dimensions. Finally, two fingerprints were obtained by looking both at the average time course:

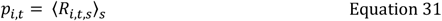

And the average response to each image:

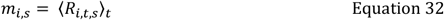

We then concatenated the vectors of elements *p*_*i,t*_ and *m*_*i,s*_ for each unit *i* to obtain its second functional fingerprint.

### Cross-correlation fingerprint

We finally computed the correlation of each unit with a reference population of units that was tracked across days. For each day *d*, we first binned the spiking activity of each unit across each half of the session using bins of 10 ms. Then, we computed the cross-correlation of each unit with every unit that was found to be tracked across days, yielding vectors *C*_*i*_of elements *c*_*i,j*_ corresponding to the instantaneous correlation coefficient of unit *i* with unit *j*. The value of the correlation of one unit with itself if the unit was part of the reference population was set to NaN. These vectors *C*_*i*_ were finally used as the third functional fingerprint.

### Fingerprint stability

The assess the similarity 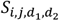 of the fingerprints of the units *i* and *j* across two days *d*_*1*_ and *d*_*2*_, we first computed the fingerprints separately for both halves of the recording sessions, yielding two fingerprints 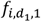 and 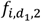 for each unit. Then, we computed the correlation of the fingerprint of units *i* and *j* across the two days and using different halves:

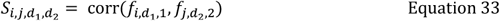

Using two different halves allowed use to compute the fingerprint’s reliability when *d*_*1*_ = *d*_*2*_.

### ROC and AUC

To quantify the amount of information present in the distributions of the correlations of the fingerprints, we computed the Receiver Operating Characteristic (ROC) curve for different populations of pairs: pairs coming from the same units, or different units within days, or pairs coming from putative matched units, or non-matched units, across days. We then computed the area under the ROC curve (AUC) to quantify this difference between distributions.

Only sessions with at least 20 matched units were taken into consideration. Moreover, in the case of the natural images responses fingerprint, these 20 units had to be reliable on the first day (test-retest reliability of the fingerprint > 0.2). For each mouse, the AUCs were then averaged across recordings locations. Similarly, the slope of AUC vs. days was computed for each recording location, and all slopes for each mouse were then averaged. Statistics were performed across animals.

#### Supplementary analyses

### Validation using alternative approaches

As a sanity check, we asked how UnitMatch performed in matching units recorded in two halves of a single recording session. These are units that were assessed to be the same across the two halves by the spike sorting algorithm (in our case, Kilosort). Consistent with the way the classifier was trained, UnitMatch tended to agree with the algorithm on these within-day matches (Figure S3A). Indeed, disagreements were rare: in 10 recordings spike-sorted individually, UnitMatch found 0.3±0.3% unexpected matches and 3.7±1.5% unexpected non-matches (Figure S3A). These might represent false positives and false negatives by UnitMatch, but they may also indicate mistakes by the spike sorting algorithm, which might have split a single unit in two, or misidentified a noisy trace or multiunit activity as a single unit.

To evaluate UnitMatch’s performance across days, we compared its output to a standard approach: running the spike sorter on the concatenated recordings^29,30^. We then ran UnitMatch directly on Kilosort’s output and compared the decisions of the spike sorter vs. UnitMatch. Running UnitMatch on the output of concatenated Kilosort yielded similar levels of unexpected matches (0.4±0.4%, N=5 mice, each two recordings) and non-matches (5.8±3.3%) within days as when the recordings were sorted separately (Figure S3A). Across days, 34.4±22.7% of units that were identified as the same unit by Kilosort were not identified as matches by UnitMatch. This is a substantial difference, but it bears remembering that spike sorting algorithms have their own biases. We thus next asked whether UnitMatch agrees with human curation.

We compared results from UnitMatch to the judgment of five expert curators. These curators were asked to judge whether pairs of waveforms belonged to the same unit (Figure S4). The pairs were shown in random order, drawn from two sets: matching pairs (as identified by UnitMatch or by the spike sorting algorithm) within and across days and an equal number of non-matching pairs. The curator was not told the identity of the pair and was not given feedback. The results show that human curators performed similarly to UnitMatch (Figure S3C-D). Both the curators and UnitMatch were generally more conservative than Kilosort (run on concatenated files) in calling a pair a match. In line with this, there was a larger overlap between matches made by UnitMatch, and pairs with stable functional scores. Especially with larger gaps between recordings, the stitched Kilosort method would overestimate the number of matched units between recordings compared to UnitMatch and stable functional scores (Figure S3E-F).

### Matching success depends on unit ‘quality’

There was quite a large variability in matching success across days, and we wanted to know to what extent this could be explained by other variables than the validity of UnitMatch. Our suspicion was that the quality of the data would be an important predictor on the number of matches. We therefore looked at the predictive value of different quality measures (Bombcell output; Fabre et al., 2023) on whether a match could be found for units included in our analysis. Area under the curve (AUC) of a receiver operating characteristic (ROC) analysis showed that indeed matching success depends on quality of the units, such as the number of (missing) spikes (Figure S5A-C), refractory period violations (Figure S5D), baseline ‘flatness’ (Figure S5E), peak amplitude (Figure S5F), and the amount of drift (Figure S5G) (AUC distribution significantly different from 0.5, t-tests, p<0.01). Surprisingly, matching success also depends on waveform duration (Figure S5H) and the number of peaks (Figure S5I), giving us direction in how to improve UnitMatch further in the future. Other quality measures, such as (Figure S5J-K) did not play a significant role (AUC not significantly different from 0.5).

## Supplementary Figures

**Figure S1.**
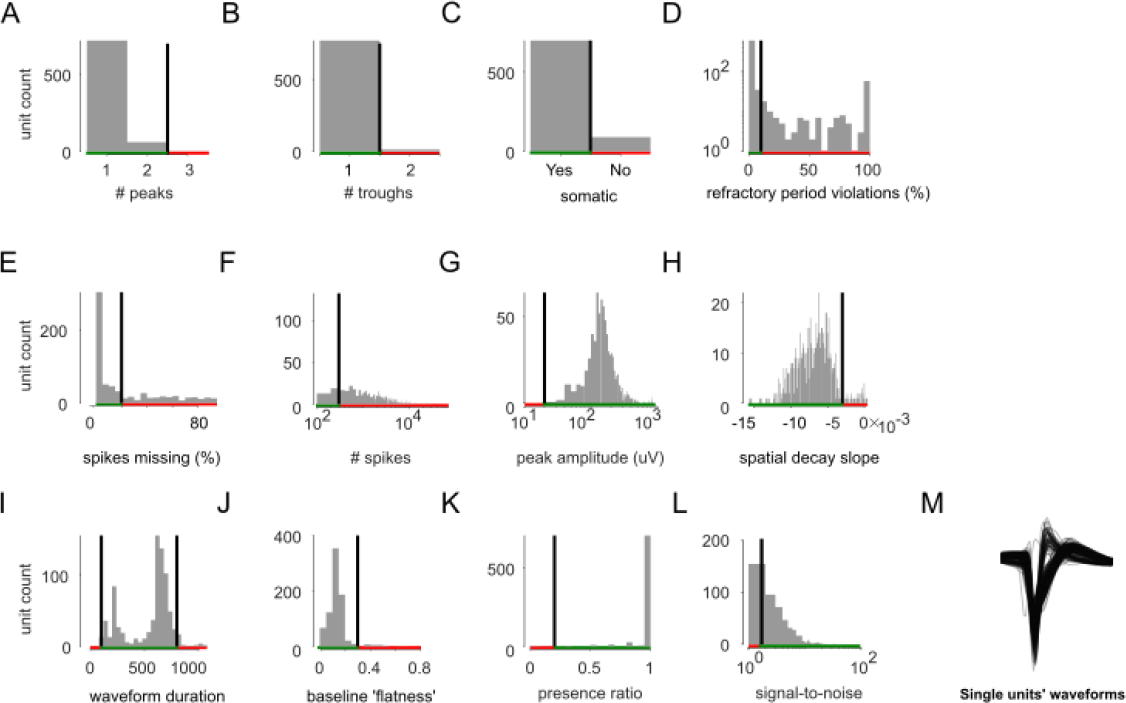
Bombcell output distributions for an example dataset. (A-L) For all the units in an example dataset (Mouse ID 1, Table S1) we measured 12 parameters. Each panel shows the number of units that passed (*green* section of the abscissa) or did not pass (*red* section) the selection based on that parameter. The parameters are: (A) Number of detected peaks. (B) Number of detected throughs. (C) Somatic waveform. (D) Estimated percentage of refractory period violations. (E) Estimated percentage of spikes below the spike sorting algorithm’s detection threshold, assuming a Gaussian distribution of spike amplitudes. (F) Total number of spikes. (G) Mean raw absolute waveform amplitude (μV). (H) Spatial decay slope (fit). (I) Waveform duration (μs). (J) Waveform baseline ‘flatness’, defined as the ratio between the maximum value in the waveform baseline and the maximum value in the waveform. (K) Presence ratio (of total recording time), defined as the fraction of bins that contain at least one spike. (L) Signal-to-noise ratio. (M) Waveforms of units that survived all quality metrics thresholds.

**Figure S2.**
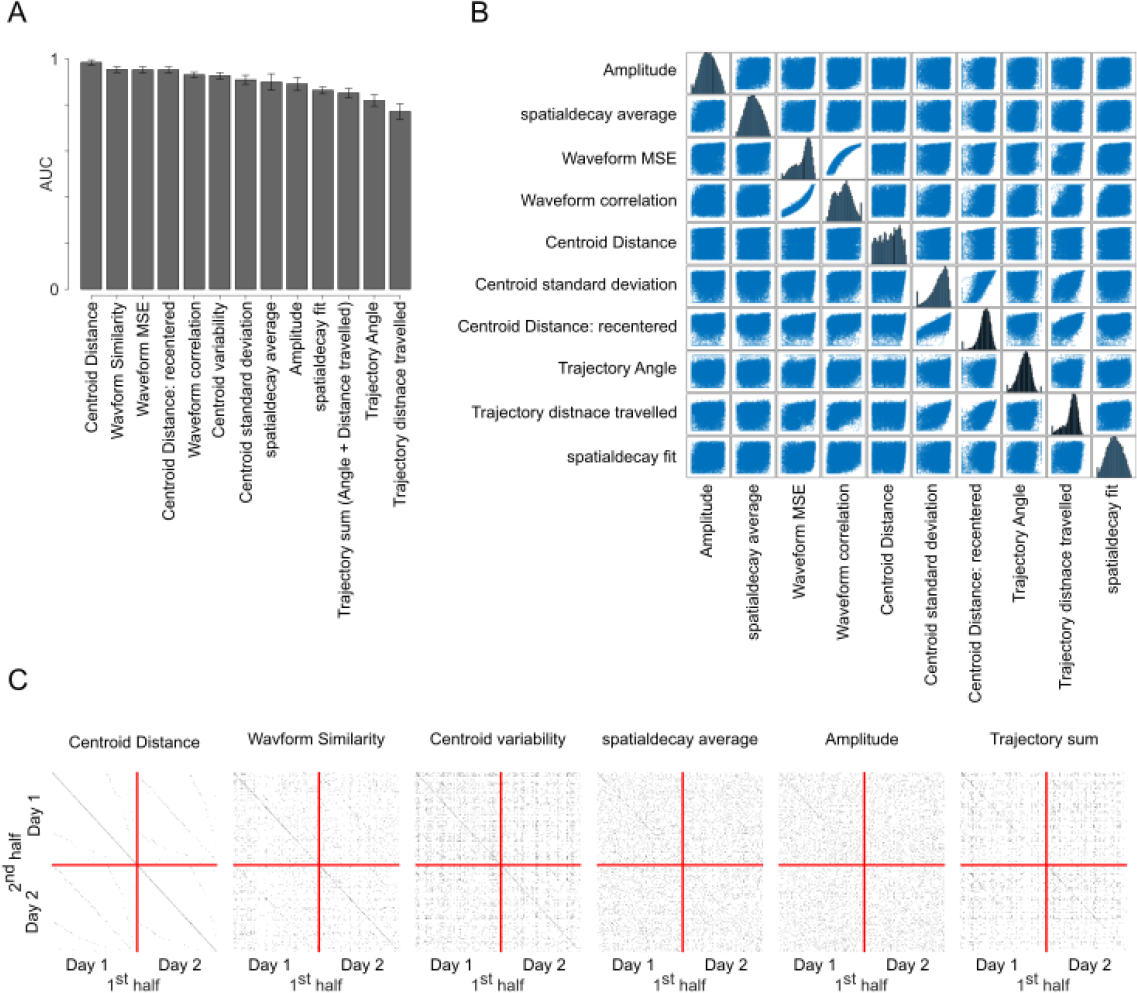
Similarity scores. (A)Area under the curve (AUC) for a receiver operating characteristic (ROC) classifying same units versus neighboring units. Large AUC values indicate that the similarity score is very informative in telling whether two waveforms come from the same unit versus whether that is not the case. (B)Cross-correlation between each pair of similarity scores. The diagonal shows histograms for the individual scores. When two parameters were very informative (large AUC scores) but correlated, we averaged them together (e.g., waveform similarity is the average of waveform MSE and waveform correlations). (C)Individual similarity scores (thresholded at same prior as total score).

**Figure S3.**
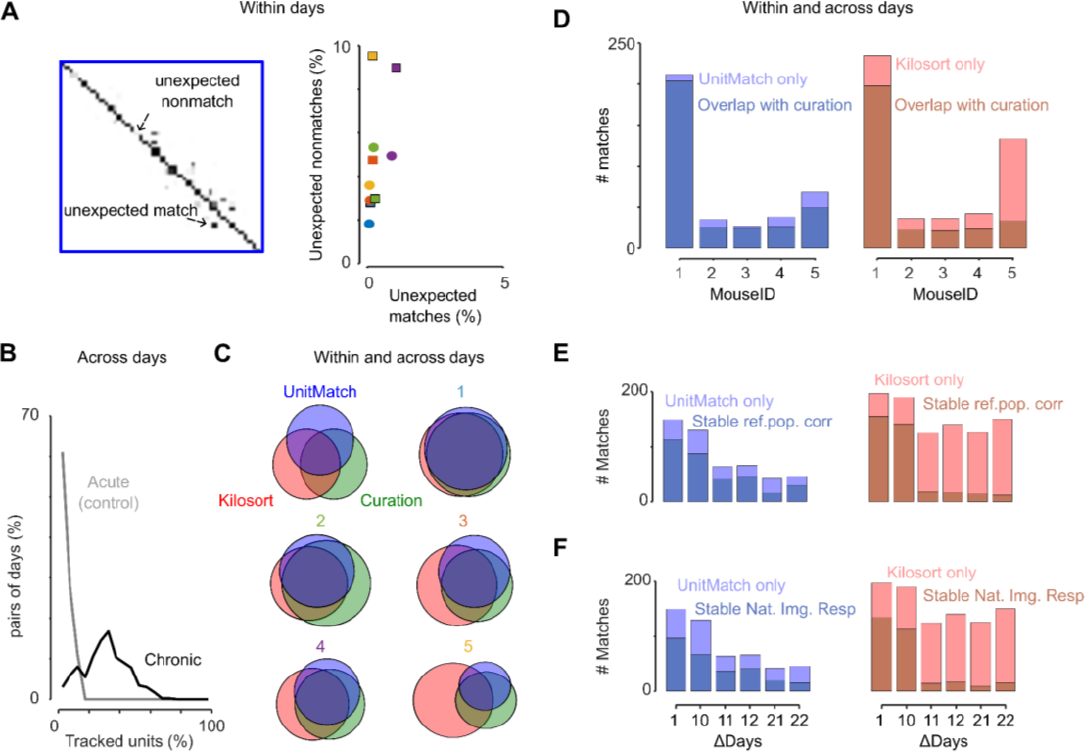
UnitMatch performance. (A)Left: Zoom in on Figure 2J on some units along the main diagonal. On the diagonal we expect p(match) to be close to 1, off diagonal we expect p(match) to be close to 0. *Right:* Unexpected matches (%) and nonmatches (%) by UnitMatch relative to units that were defined as good single units within a day by Kilosort. UnitMatch was either run on individually sorted data (*circles*), or on concatenated data (*squares*). Colors depict individual mice, and refer to colors in (B)Pairs of consecutive recording days (%) for acute (gray) and chronic (black) recordings as a function of the percentage of tracked units. The percentage of tracked units is defined as the number of matched units between two consecutive recording days divided by the number of units on the recording day with the least recorded units. (C)Venn diagrams for five individual mice illustrating the overlapping pairs of units assigned as ‘match’. For some data sets all three methods (Kilosort, UnitMatch, and curation) largely agreed, whereas for other datasets the three methods did not agree. In that case, Kilosort typically assigned more pairs as a match than the other two methods. (D)Units matched within or across two (concatenated) consecutive days by UnitMatch (left) and Kilosort (right) for five mice. Dark parts of the bars show the overlap with curated matches, light parts of the bar the matches additionally made by the respective algorithm. (E)Number of units matched across many (concatenated) days (x-axis) by UnitMatch (left) and Kilosort (right). The overlap with stable reference population correlations is shown (most stable pairs only). Four recordings were stitched together for one mouse (ID 1). (F)same as (E) but showing the overlap with stable natural image responses (most stable pairs only). Note that in (E) and (F) the criteria for a unit to be labeled as “stable” are highly stringent (a unit’s fingerprint on day 1 must be best correlated with its fingerprint on day n, above all other units).

**Figure S4.**
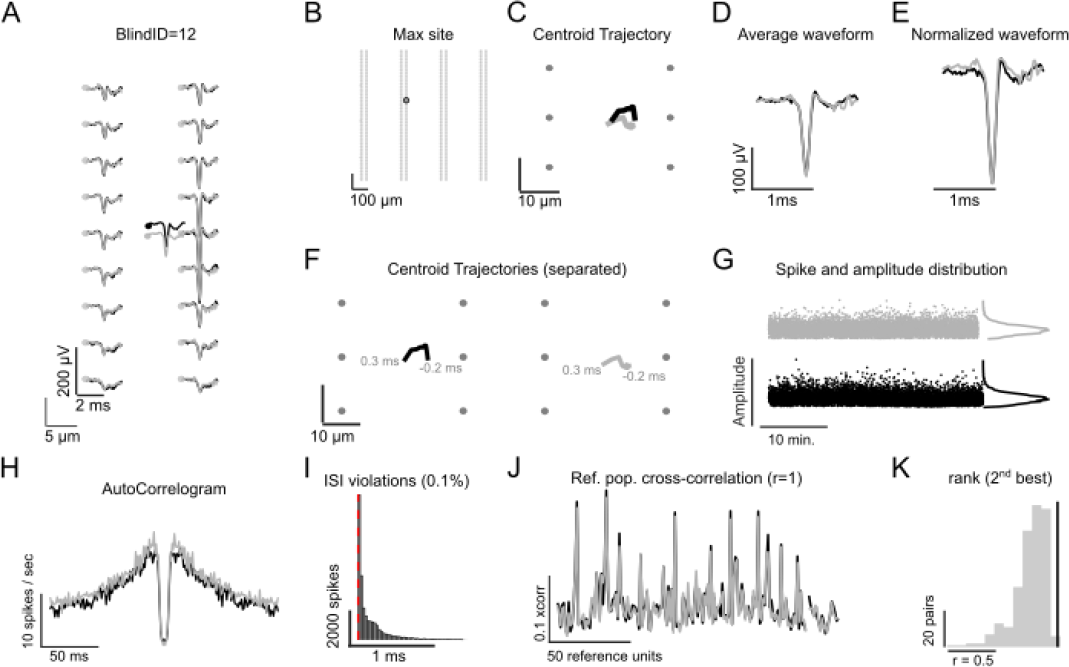
Expert curation. Five expert curators were asked to judge whether two waveforms came from the same unit using a figure like this. Example BlindID 12: ‘black’ and ‘gray’ unit. (A)Average waveform across recording sites. (B)Maximum recording site indicated (note, they overlap) on a 4-shank Neuropixels. (C)Centroid trajectories. (D)Average waveforms. (E)Normalized waveforms (peak to base stretching). (F)Same as (C) but shown next to each other for blue and cyan unit. (G)Spike times (x-axis) versus amplitude (y-axis), with the amplitude distribution next to it. Note that the amplitudes for ‘gray’ are drawn above ‘black’ for visibility. (H)Autocorrelogram. (I)Inter spike interval distribution. (J)Reference population cross-correlation, which was 1 for this specific pair. (K)reference population cross-correlation values between this pair of units (black line), relative to other possible pairs of units (distribution). Rank 2 means this cross-correlation value is the second highest of all possible pairs.

**Figure S5.**
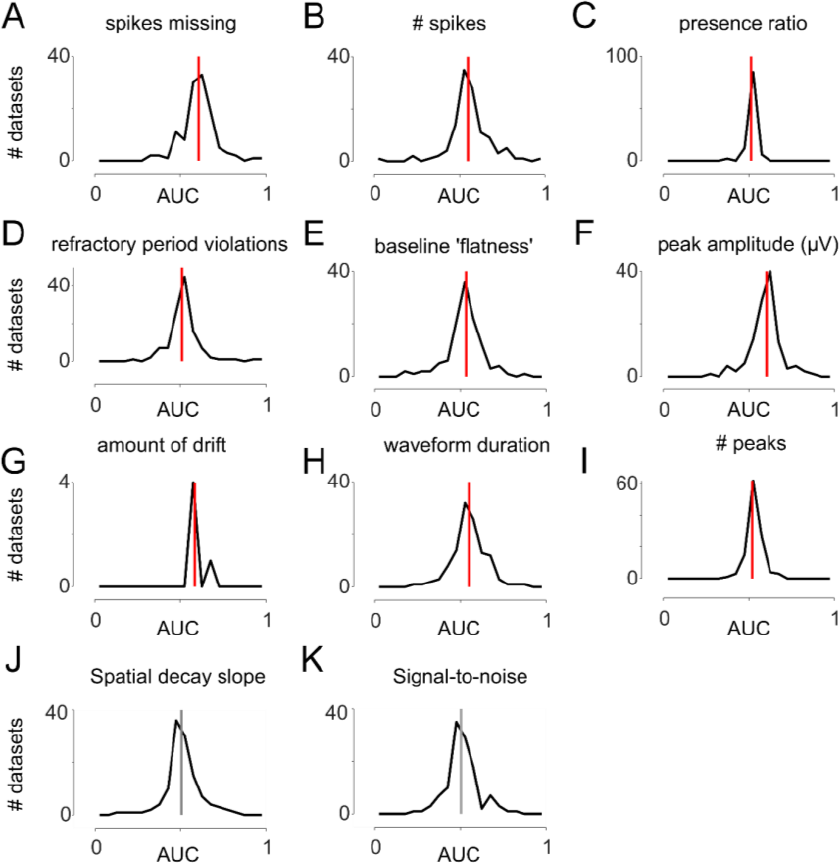
Quality metrics significantly predict whether UnitMatch can find a match for a unit. An AUC can be computed for each quality metric to quantify whether this metric differs across matches and non-matches. Each plot shows the distribution of AUC values and the median AUC value, across all datasets. (A) spikes missing, (B) number of spikes, (C) presence ratio, (D) number of refractory period violations, (E) baseline ‘flatness’, (F) peak amplitude, (G) amount of drift, (H) waveform duration, (I) number of peaks, (J) spatial decay slope, and (K) signal-to-noise ratio. Distributions for A-I were significantly different from 0.5 (t-test, p<0.01), suggesting these quality metrics have predictive power on whether a unit is likely to be matched with another unit. Distributions for J and K were not significantly different from 0.5.

**Table S1.** Information on animals, implants, and software used. For every animal (ID) we specify what type of implant was used, what probe type of Neuropixels (Npx) probe was used and how many, and which areas the implant reached. We also specify the Kilosort algorithm used for preprocessing. pyKS: **https://github.com/MouseLand/pykilosort**. We used the ‘develop’ branch. KS2: **https://github.com/MouseLand/Kilosort/releases/tag/v2.0**, which was first preprocessed using CatGT: **https://github.com/billkarsh/CatGT**. Abbreviations: V1: primary visual cortex, HC: hippocampus, DMS: dorsomedial striatum, RSP: retrosplenial cortex, SC: superior colliculus, THAL: thalamus, FC: frontal cortex, STR: striatum. Mouse ID 1 was used for all example figures (Ex.) including Fig. 4A-D,F-I,K-N. All mice in Fig. 4E,O were also used in figure S5. Note: mouse 2 and 19 are the same mouse, preprocessed differently.

| ID | IMPLANT | PROBE TYPE | REGIONS | KILOSORT | EX. | FIG 4E,O | FIG 4J | FIG 5 | S3 |
| --- | --- | --- | --- | --- | --- | --- | --- | --- | --- |
| 1 | Cemented | Npx 2.0 (4s) | V1, HC | pyKS | x | x | x |  | x |
| 2 | Ultralight | Npx 1.0 | DMS | pyKS |  |  |  |  | x |
| 3 | Apollo | Npx 2.0 (4s) x2 | RSP, SC | pyKS |  | x |  |  | x |
| 4 | Apollo | Npx 2.0 (4s) | V1, HC, THAL | pyKS |  | x |  |  | x |
| 5 | Apollo | Npx 2.0 (4s) | V1, HC | pyKS |  | x | x |  | x |
| 6 | Cemented | Npx 1.0 | V1, HC | pyKS |  | x | x |  |  |
| 7 | Cemented | Npx 2.0 (1s) | V1, HC | pyKS |  | x |  |  |  |
| 8 | Cemented | Npx 2.0 (4s) | V1, HC | pyKS |  | x | x |  |  |
| 9 | Apollo | Npx 2.0 (4s) x2 | FR, STR | pyKS |  | x |  |  |  |
| 10 | Apollo | Npx 2.0 (4s) x2 | FR, STR | pyKS |  | x |  |  |  |
| 11 | Isogai | Npx 1.0 | V1, HC | pyKS |  | x | x |  |  |
| 12 | Isogai | Npx 1.0 | V1, HC | pyKS |  | x | x |  |  |
| 13 | Isogai | Npx 1.0 | V1, HC | pyKS |  | x |  |  |  |
| 14 | Haesler | Npx 1.0 | V1, HC | pyKS |  | x | x |  |  |
| 15 | Apollo | Npx 2.0 (4s) | V1, HC | pyKS |  | x | x |  |  |
| 16 | Apollo | Npx 2.0 (4s) | V1, HC | pyKS |  | x | x |  |  |
| 17 | Apollo | Npx 2.0 (4s) | V1, HC | pyKS |  | x |  |  |  |
| 18 | Apollo | Npx 2.0 (4s) | V1, HC | pyKS |  | x | x |  |  |
| 19 | Ultralight | Npx 1.0 | DMS | KS2 |  |  |  | x |  |

